# Quantitative PCR assays to detect humpback whale (*Megaptera novaeangliae*), shortbelly rockfish (*Sebastes jordani*), and common murre (*Uria aalge*) in marine water samples

**DOI:** 10.1101/2020.08.17.240275

**Authors:** Elizabeth A. Andruszkiewicz, Kevan M. Yamahara, Collin J. Closek, Alexandria B. Boehm

## Abstract

Monitoring aquatic species by identification of environmental DNA (eDNA) is becoming more common. In order to obtain quantitative datasets for individual species, species-specific quantitative PCR (qPCR) assays are required. Here, we present detailed methodology of qPCR assay design and testing, including *in silico, in vitro*, and *in vivo* testing, and comment on the challenges associated with assay design and performance. We use the presented methodology to design assays for three important marine organisms common in the California Current Ecosystem (CCE): humpback whale (*Megaptera novaeangliae*), shortbelly rockfish (*Sebastes jordani*), and common murre (*Uria aalge*). All three assays have excellent sensitivity and high efficiencies ranging from 92% to 99%. However, specificities of the assays varied from species-specific in the case of common murre to the genus-specific shortbelly rockfish assay, to the humpback whale assay which cross-amplified with other two other whale species, including one in a different family. All assays detected their associated targets in complex environmental water samples.

## Introduction

Researchers across the globe have started monitoring aquatic species by identifying environmental DNA (eDNA) captured in water samples [1]. All organisms are constantly shedding DNA, which remains in an environment (i.e., soil, air, water), hence the term “environmental DNA,” or “eDNA” [1,2]. Monitoring by eDNA has several advantages over traditional monitoring methods such as trawl nets or visual identification. eDNA sampling is non-invasive and is independent of visually identifying the organism, thus the method is not subject to avoidance or identification error [3]. Small volume (i.e., 1 L) water samples can be analyzed for multiple taxonomic groups by applying molecular methods such as quantitative PCR (qPCR) or metabarcoding [4,5]. Major limitations of eDNA methods include uncertainty about the organism number or biomass present based on the eDNA concentration and the exact location and time of eDNA shedding from organisms [6–8]. However, the interest in using eDNA to identify marine organisms is growing and more assays are needed to identify important species.

Although results using eDNA metabarcoding provide information about many species from a single water sample, studies have demonstrated that the resulting data are not necessarily quantitative [9,10]. Furthermore, the metabarcoding method is prone to false negatives due to various technical challenges, including different target species having different affinities to PCR primers [7,9]. qPCR methods may be less prone to false negatives and studies have shown correlation between biomass and eDNA concentration [6,11,12]. The difficulty in using qPCR assays is that for every species of interest, a new assay must be designed and tested to evaluate its performance. The design of a qPCR assay depends upon having reliable genetic information from many individuals of the target species and finding a short region of a gene that will be specific to the target species and also suitable for the reaction chemistry of qPCR to ensure the assay is efficient. Finally, testing must be performed on other non-target species that are either co-occurring or closely related, as well as testing with water samples known to contain target DNA to evaluate performance in a complex matrix. Thus, the process can be laborious and time intensive. However, once the assay is developed, its use for biomonitoring has great potential.

We focus on three species of particular interest in the California Current Ecosystem (CCE): humpback whale (*Megaptera novaengliae*), shortbelly rockfish (*Sebastes jordani*), and common murre (*Uria aalge*). The CCE is a productive eastern boundary upwelling system along the west coast of the United States from Washington, USA to Baja California [13,14]. The CCE is well studied and particularly important to monitor due to its high primary productivity as a result of the upwelling system. The productivity in turn makes the CCE economically valuable. Therefore, it is necessary to monitor populations of taxa and biodiversity for conservation efforts. The CCE includes many protected areas, including federal National Marine Sanctuaries, state Marine Conservation Areas and Reserves, and local conservation areas.

Within the food web of the CCE, humpback whales, shortbelly rockfish, and common murre are three important organisms [15]. Humpback whales are protected under the federal Marine Mammal Protection Act enacted in 1972 when the species was close to extinction. Humpback whales are tertiary consumers and are frequently found in Monterey Bay due to the abundance of small fish and zooplankton [16,17]. Monitoring of humpback whale is traditionally carried out using visual surveys and can be sparse in space and time [18]. Recently, the use of citizens to conduct humpback whale surveys has gained momentum through citizen science campaigns [19].

Rockfish are secondary consumers in the CCE food web, consuming plankton and small fish [15]. The National Oceanic and Atmospheric Administration (NOAA) currently monitors populations of rockfish, including shortbelly rockfish, during its annual Rockfish Recruitment and Ecosystem Assessment Survey (RREAS) in the central CCE [15,20,21]. The RREAS has been conducted since 1983, and represents one of the longest time series of epipelagic juvenile fishes [22]. Shortbelly rockfish are one of over 100 species in the genus, *Sebastes*. Despite the long-term surveys, it can be difficult to make species assignments to juveniles that do not yet have identifying features of adults [23]. Net trawl surveys results in mortality of the captured organisms and can even result in mortality of individuals that escape the trawl net [24].

Common murre are piscivorous, top predator seabirds, that are abundant in the CCE. Their seasonal abundance has been correlated with availability of juvenile rockfish which can be a main source of prey [25,26]. Seabirds are monitored annually during the RREAS by visual survey (line transect), where an observer uses pre-determined distance intervals to estimate densities of seabirds [18,21]. This method is reliant on human observation and thus is subject to misidentification or false negatives [27].

The reliance on visual observations and trawling for abundance assessments of these three organisms gives rise to spatially and temporally sparse abundance data sets. The goal of this study is to develop eDNA focused methods to identify these organisms from water samples. We therefore designed and tested hydrolysis probe-based quantitative PCR (qPCR) assays for humpback whale (*Megaptera novaeangliae*), shortbelly rockfish (*Sebastes jordani*), and common murre (*Uria aalge*). The assays were designed and tested *in silico*, *in vitro* in the laboratory, and *in vivo* using field samples suspected to contain eDNA from the organisms. Future efforts to test ecological hypotheses or carry out biomonitoring of these three marine species can utilize these assays. Finally, we include discussion of the challenges associated with qPCR assay design and suggestions for addressing these challenges.

## Materials and methods

### qPCR assay design

Assays were designed for the three organisms using the Geneious software (version 11.1.5), which uses a modified version of Primer3 (Version 2.3.7) [28]. For each organism, all sequences from the BLAST nucleotide (nt) database obtained from tissue vouchers were downloaded for a target gene (search completed on 16 August 2018). For humpback whale, the assay targeted the d-loop control region of the mitogenome. We limited the downloaded sequences for humpback whale assay design to include only those sequences obtained from organisms from the North Pacific (n = 180, S1 Table). The common murre and shortbelly rockfish assays both target the mitochondrial cytochrome c oxidase subunit I (COI) gene. These genes were chosen based on the availability of sequences for target species in the nt database. There were 16 common murre COI gene sequences and 5 shortbelly rockfish gene sequences at the time of our search, so all were downloaded for assay design (S1 Table).

Downloaded sequences were aligned using the default parameters in Geneious for a MUSCLE alignment and a consensus sequence was developed for each target. For all three targets, the assessment showed pairwise percent identity >97%. Thus, a representative sequence was chosen for each target to use for primer design from the National Center for Biotechnology Information (NCBI) database (humpback whale: GenBank Accession GQ353077, shortbelly rockfish: JQ354411, common murre: GU572157) (S1 Table). The “Design new primers” tool in Geneious was used in conjunction with the criteria outlined in Table 1, which included setting limits on product size, primer/probe size, primer/probe melting temperature (T_m_), and primer/probe GC content [29–32]. Ten primer/probe sets were returned by the software and were evaluated manually using the criteria outlined in Table 1, including optimizing the ΔT_m_ between primers and probe, no continuous occurrence of 4 or more of the same base, avoiding a thymine base at the 3’ end of the primer, including guanine or cytosine bases at the end of the primers, and no guanine at the 5’ end of the probe to reduce self-quenching of the fluorophore.

**Table 1.**
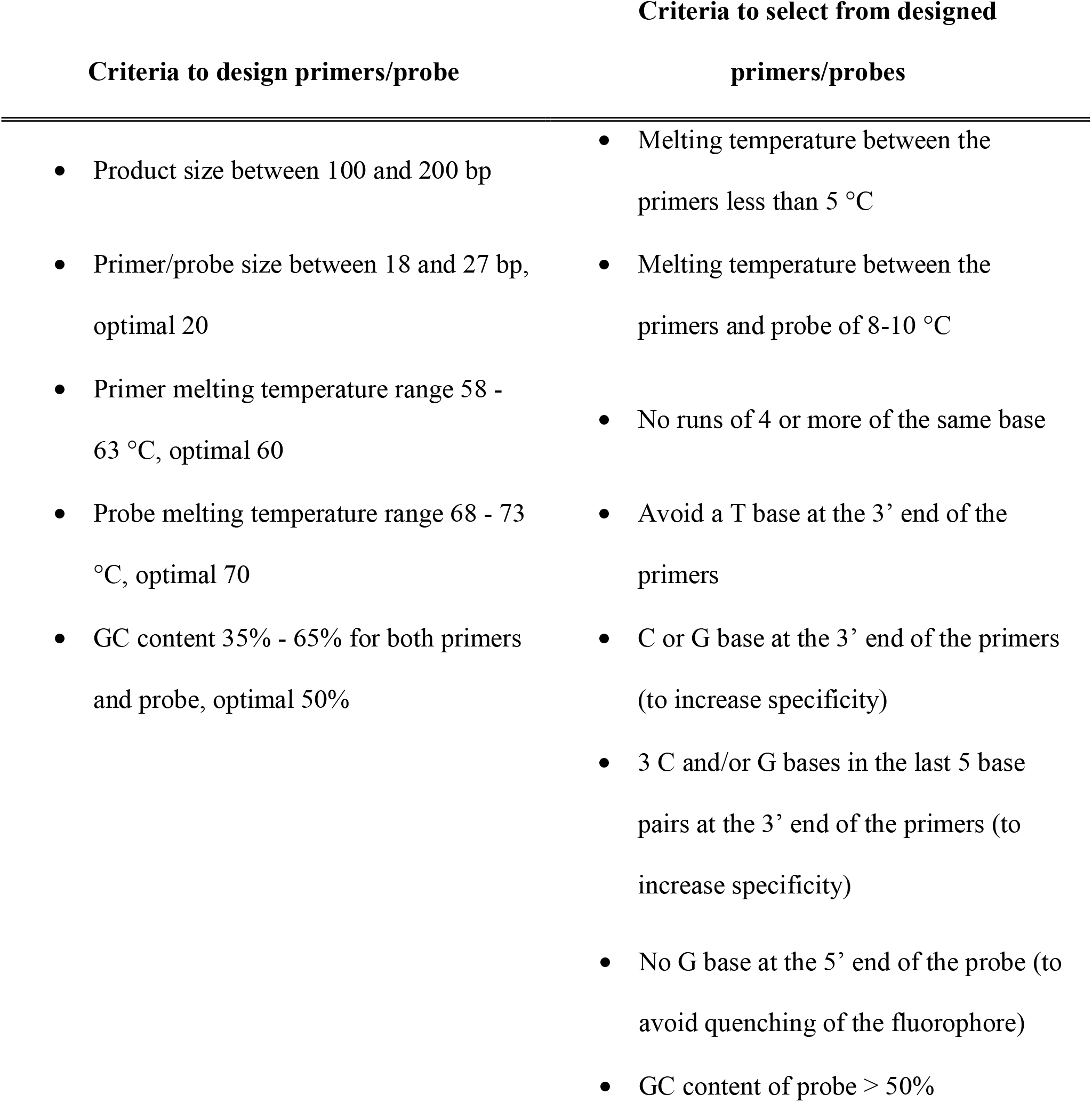
Criteria used for designing primer/probe sets and criteria used to assess primer/probe sets in silico.

The primer and probe set for each assay that adhered to the greatest number of criteria were subsequently used to test specificity *in silico* and sensitivity and specificity *in vitro*. These primer and probe sets are hereafter referred to as the “preliminary primer/probe sets”.

### *In silico* specificity testing

*In silico* testing was performed using the Geneious software “Blast” tool for the preliminary primer/probe sets using the default parameters. The primer/probe sequences were defined as the “Query” and the search was performed using the entire nucleotide collection (nt/nr) with the default parameters, except that the “maximum hits” criterion was set to 9,000. For each assay, database hits obtained for the forward primer, probe, and reverse primer were combined to identify entries in the nt database that contained sequences that matched both primers and probe sequences (search completed on 30 March 2019, S2 Table).

We generated sequence logo plots to illustrate the *in silico* specificity of the two primers and probe for each assay. Sequence logo plots were generated using the R package “ggseqlogo” [33] to illustrate the conserved and non-conserved nucleotide bases in the primer and probe binding regions [34]. Plots show (1) the relative frequency of base pair occurrence of the primers and probe by aligning all of the sequences for target species for each assay and (2) the relative frequency of base pair occurrence of the primers and probes using an alignment of non-target taxa. The latter was generated by aligning the regions of the forward/reverse primers and probe from all other non-target taxa within the same family of the target individual (e.g., all species within the family Balaenopteridae for humpback whale).

### Library of target and non-target samples for *in vitro* testing

In order to test and optimize each assay, we obtained tissue samples of individuals from both target species and non-target species to determine the sensitivity and specificity of each assay, respectively. We obtained tissue samples from 4 humpback whales, 3 shortbelly rockfish, and 3 common murre individuals (S3 Table). The humpback individuals were skin biopsies provided from either NOAA SWFSC (Permit # SR395) or The Marine Mammal Center (TMMC) (Permit # 19091). The shortbelly rockfish individuals were from fin clips or tissue provided by NOAA Southwest Fisheries Science Center (SWFSC) (no permit required). The murre individuals were provided by the California Department of Fish and Wildlife in Santa Cruz, California, USA in the form of liver tissue samples (no permit required).

Non-target species samples were obtained as tissue or archived tissue DNA extracts for *in vitro* specificity testing (S3 Table). The humpback whale assay was tested against a total of 10 other species (n = 3 for 8 species, n = 1 for 2 species), including 5 other whale species (blue whale, fin whale, minke whale, grey whale, bowhead whale). The shortbelly rockfish assay was tested against 23 other species (n = 3 for 7 species, n = 1 for 16 species), including 12 different species within the rockfish genus, *Sebastes*. The common murre assay was tested against DNA from 7 other birds (n = 1 individual for each of 5 species, n = 2 individuals for each of 2 species, S3 Table). S3 Table provides metadata on the samples used for specificity testing and permit information for marine mammals.

For non-target samples that were provided as tissue, DNA was extracted in our laboratory from each sample either using Qiagen DNeasy Blood and Tissue Kit (Qiagen, Valencia, CA) or by salting out [35] and the DNA extract was quantified using Qubit dsDNA HS assay (Life Technologies, Grand Island, NY). Samples that were provided as DNA extract were quantified using Qubit dsDNA HS assay (Life Technologies, Grand Island, NY).

### Assay optimization

Using one individual of each target species from our library, assays were optimized by varying primer/probe concentrations and annealing temperatures to achieve sensitive and high efficiency assays. For each assay, two non-target samples were included to check for specificity during assay optimization. Primer and probe concentrations tested ranged from 0.2 – 0.8 μM for primers and 0.1 – 0.2 μM for probes for a total of 6 combinations of primer/probe concentrations (primer/probe: 0.2/0.1, 0.2/0.2, 0.5/0.1, 0.5/0.2, 0.8/0.1 μM). The following reaction chemistry was used for assay development: Taqman Universal Mastermix II (1x), forward and reverse primer (varying concentration), probe (varying concentration), 2 μL of DNA extract, and molecular-biology-grade water (Sigma-Aldrich, St. Louis, MO).

Annealing temperatures tested ranged from 60 – 64°C in 1°C increments. The cycling parameters were the same for all of the assays and were as follows: were 95°C for 5 min followed by 40 cycles of 95°C for 15 s, the annealing temperature being tested for 30 s and 72°C for 30 s. The C_q_ (quantification cycle) threshold was set at either 0.01 (humpback whale) or 0.02 (murre and shortbelly rockfish). No template controls (NTCs) were added with each plate tested using molecular grade water in lieu of DNA extract. All experiments were performed using a StepOne Plus Real-Time PCR System (Applied Biosystems, Foster City, CA).

Assays were also optimized to improve specificity by adding bovine serum album (BSA) if needed. For each assay, specificity, sensitivity, and efficiency were evaluated to determine the optimal primer and probe concentrations, annealing temperature, and the addition of BSA. After assay optimization, the preliminary primer/probe sets were all considered to be the final primer/probe sets and their sensitivity and specificity were determined before applying the assays to environmental samples.

### *In vitro* sensitivity and specificity testing

For the *in vitro* sensitivity testing, each primer/probe set was tested against a dilution series using DNA extracted from all individuals of each target species as template (n = 4 humpback whales, n = 3 shortbelly rockfish, n = 3 common murres). The highest concentration was 200 pg DNA per reaction and the lowest was 2 fg per reaction; 1:10 dilutions of the highest concentration was used for intermediate dilutions. The dilution series was tested using triplicate reactions.

The *in vitro* specificity testing was conducted using all non-target samples from the library by including triplicate reactions of each sample, where 1-2 ng of DNA was added per reaction. This test was conducted using the final primer/probe sets and cycling parameters after optimization.

### Data analysis

We identified the limit of quantification (“LOQ”) for each assay as the lowest concentration of target DNA for which all three triplicates were consistently assigned a cycle quantification (C_q_) value. Samples assigned a C_q_ value lower than that of the LOQ were considered positive. Samples that were not assigned a C_q_ value were considered non-detects (“ND”) and considered negative. No sample amplified at a C_q_ value higher than the that of the LOQ so there was no need to consider how to categorize measurements in this range.

Sensitivity was calculated as the number of positive target samples divided by the total number of target samples. It is the ratio of true positives to the sum of false negative and true positives. Specificity was calculated by dividing the number of true negatives by the sum of the false positives and true negatives.

### Application *in vivo*

Environmental water samples for *in vivo* testing were collected from Monterey Bay, California, USA and the Monterey Bay Aquarium Diving Birds Exhibit. Samples for testing the humpback whale assay were collected from the surface of the water column using a 10 L 10% HCl-acid washed, autoclaved polypropylene carboy (Nalgene, Rochester, NY) aboard the *R/V Paragon*. The samples used to test the shortbelly rockfish assay were collected aboard the *R/V Reuben Lasker* at 40 m depth using a Niskin array during conductivity-temperature-depth (CTD) casts. One liter water samples were collected from the Diving Birds Exhibit at the Monterey Bay Aquarium for testing the common murre assay using a 10% HCl-acid washed bottle. Water samples collected from the *R/V Paragon* were filtered and preserved with the Environmental Sample Processor using 0.22 μm pore size 255 mm diameter durapore filters (Millipore, Burlington, MA) [36]. All other water samples were filtered through 0.22 μm pore size 47 mm diameter durapore filters using DNA-clean, sterilized vacuum filtration devices. Filters were stored at −80°C until DNA extraction (within 16 months). DNA was extracted from filters using a previously described Qiagen DNeasy Tissue Extraction with bead-beating [37]. Aliquots of DNA were stored at −80°C until used as template in qPCR reactions (within 11 months).

Amplification of the field samples by qPCR was performed using 20 μL reactions with 2 μL of template using the reaction chemistry and cycling parameters of the optimized assays. All environmental samples were diluted 1:5 to reduce the chance of PCR inhibition. Standard curves for each assay were generated using serial dilutions of DNA from one target individual in order to quantify DNA in the environmental samples. Six ten-fold or five-fold dilutions were run in triplicate in each qPCR plate starting with 200 pg per reaction and ending with 2 fg per reaction for ten-fold dilutions or 64 fg per reaction for five-fold dilutions. Standard curve data were used to create a regression of DNA concentration per reaction versus C_q_ to calculate concentrations of unknown samples.

## Results

### Humpback whale assay final design and performance

The humpback whale assay targets the d-loop control region of the mitogenome. The primers/probe sequences are: F 5’ GCCGCTCCATTAGATCACGA 3’, R 5’ TGGCCCTGAAGTAAGAACCAG 3’, P 5’ FAM - TCGCACCGGGCCCATCAATCGT - BHQ 3’ (Table 2). After optimizing primer/probe concentrations and annealing temperature, reactions for amplification are 20 μL in volume with the following: Taqman Universal Mastermix II (1x), 0.2 mg/ml bovine serum album (BSA), forward and reverse primer (0.2 μM), probe (0.1 μM), 2 μL of DNA extract, and molecular-biology-grade water (Sigma-Aldrich, St. Louis, MO). The cycling parameters were as follows: were 95°C for 5 min followed by 40 cycles of 95°C for 15 s, 60°C for 30 s and 72°C for 30 s.

**Table 2.**
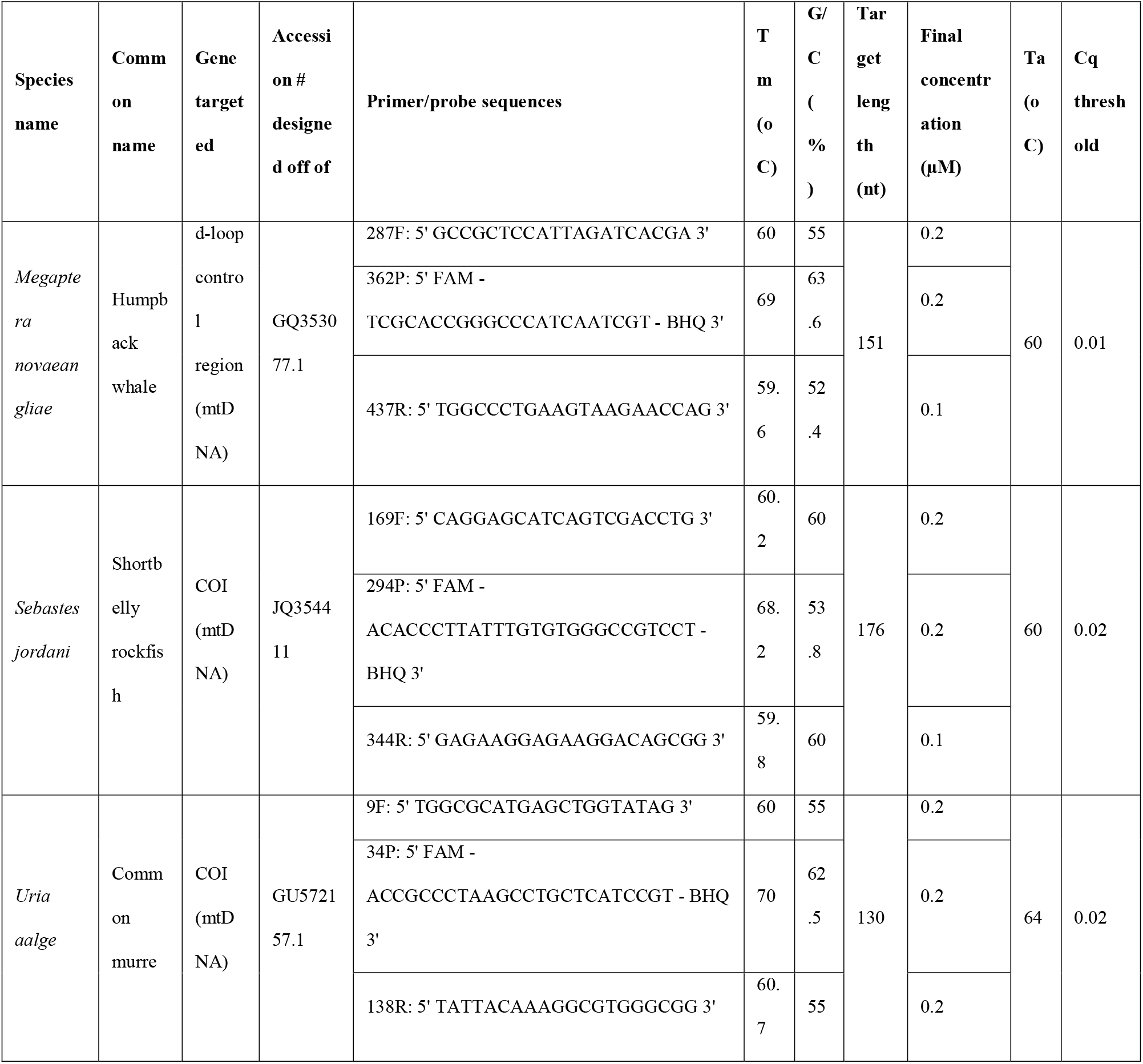
Genus-specific primers and probes designed. Target organism, primer and probe sequences, gene target, fragment size, final concentrations of primers and probe, slope, intercept, limit of quantification, and efficiency of assay.

During *in vitro* sensitivity testing, DNA from all four humpback whale individuals amplified, and thus the sensitivity of the assay was 100% (Table 3). These results match the *in silico* results demonstrated by the logo plot showing that target sequences have the exact same sequences in the binding region of the forward primer, reverse primer, and probe (i.e., all base pair relative frequencies are 1 at each position; Fig 1, Panel A, top plots).

**Fig 1.**
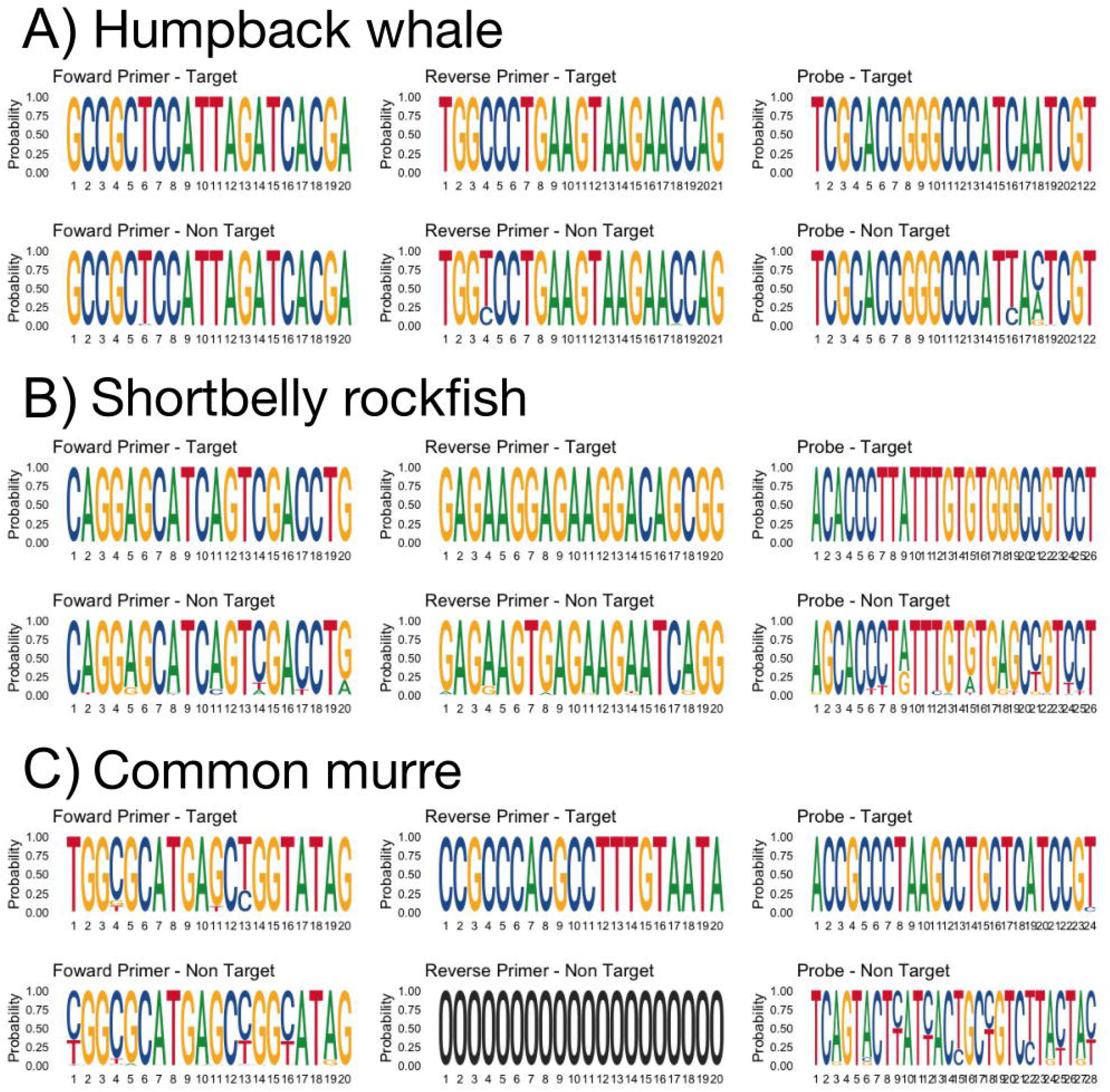
Sequence logo plots of *in silico* sensitivity and specificity of primers and probes for each qPCR assay. Panel A shows humpback whale, Panel B shows shortbelly rockfish, and Panel C shows common murre. For each panel, the top plots are generated by aligning the regions of the forward primer, reverse primer, and probe, respectively, of target species for each assay and the bottom plots are generated by aligning the same regions for non-target species. The x-axis is the order of base pairs in the primer or probe and the y-axis is the relative frequency of base pair occurrence, with the size of each letter corresponding to its relative frequency. Letters are stacked if multiple base pairs occur at that position in the primer or probe and relative frequencies always sum to 1. The reverse primer for common murre shows as “0” for all base pairs because no contigs were found when non-target species (within the family Alcidae) were aligned with the reverse primer sequence. All sequences are shown from 5’ to 3’.

**Table 3.** Sensitivity and specificity of each assay.

|  | Humpback<br>whale | Shortbelly<br>rockfish | Common<br>murre |
| --- | --- | --- | --- |
| Total target individuals | 4 | 3 | 3 |
| True positives (TP) | 4 | 3 | 3 |
| False negatives (FN) | 0 | 0 | 0 |
| Total non-target individuals | 26 | 37 | 9 |
| True negatives (TN) | 21 | 29 | 9 |
| False positives (FP) | 5 | 8 | 0 |
| Sensitivity: TP/(FN+TP) | 100% | 100% | 100% |
| Specificity: TN/(FP+TN) | 81% | 78% | 100% |

The specificity testing of the assay *in silico* returned matches to 4 additional species besides humpback whale: *Balaena mysticetus* (bowhead whale), *Balaenoptera acutorostrata* (minke whale), *Eubalaena australis* (Southern right whale), and *Eubalaena glacialis* (North Atlantic right whale) (S2 Table). *In vitro* testing included minke whale and bowhead whale (from the previous list), as well as blue whale, fin whale, and grey whale (Table 4), but did not include bowhead or Southern right whale. More generally, the *in silico* specificity illustrated in the logo plots indicated potential for cross-amplification from other taxa within the family, demonstrated by the relatively few differences in base pair frequencies in the assay between target and non-targets (1/20 in forward primer at low frequency, 2/21 in reverse primer at low frequency, and 2/22 in the probe; Fig 1, Panel A, bottom plots).

**Table 4.** Results of *in vivo* specificity testing. The three target organisms are shown on the right. Grey shading indicates that the assay was not tested for specificity using the organism on in the first column. “NA” indicates no amplification (i.e., assay is specific). For the non-target species that amplified, the C_q_ values are shown for each individual. 1-2 ng of DNA was included in each reaction; see Fig 2 for C_q_ values for target individuals.

| Specificity Testing |  |  | Target Organisms |  |  |
| --- | --- | --- | --- | --- | --- |
| <u>Species name</u> | <u>Common name</u> | <u>Individuals</u> | *Megaptera novaeangliae* | *Sebastes jordani* | *Uria aalge* |
| *Uria lomvia* | Thick-billed murre | n=1 |  |  | NA |
| *Cephus grille* | Black guillemot | n=1 |  |  | NA |
| *Alle alle* | Little auk | n=2 |  |  | NA |
| *Larus pacificus* | Pacific gull | n=1 |  |  | NA |
| *Larus delawarensis* | Ring-billed gull | n=1 |  |  | NA |
| *Larus heermanni* | Heermann's gull | n=1 |  |  | NA |
| *Sebastes auriculatus* | Brown rockfish | n=1 |  | NA |  |
| *Sebastes carnatus* | Gopher rockfish | n=1 |  | NA |  |
| *Sebastes caurinus* | Copper rockfish | n=1 |  | NA |  |
| *Sebastes entomelas* | Widow rockfish | n=3 |  | NA |  |
| *Sebastes flavidus* | Yellowtail rockfish | n=3 |  | NA |  |
| *Sebastes goodei* | Chilipepper rockfish | n=3 |  | NA |  |
| *Sebastes hopkinsi* | Squarespot rockfish | n=3 |  | 24.8, 24.2, NA |  |
| *Sebastes miniatus* | Vermillion rockfish | n=1 |  | NA |  |
| *Sebastes nebulosus* | China rockfish | n=1 |  | NA |  |
| *Sebastes paucispinis* | Bocaccio rockfish | n=3 |  | NA |  |
| *Sebastes saxicola* | Stripetail rockfish | n=3 |  | 25.6, 24.8, 24.5 |  |
| *Sebastes semicinctus* | Halfbanded rockfish | n=3 |  | 23.7, 23.2, 22.9 |  |
| *Cottus spp.* | Sculpin | n=1 |  | NA |  |
| *Ophiodon elongatus* | Lingcod | n=1 |  | NA |  |
| *Sardinops sagax* | Pacific sardine | n=1 |  | NA |  |
| *Scomber japonicus* | Pacific chub mackerel | n=1 |  | NA |  |
| *Engraulis mordax* | Northern anchovy | n=1 |  | NA |  |
| *Clupea pallasii* | Pacific herring | n=1 |  | NA |  |
| *Seriola lalandi* | Yellowtail jack | n=1 |  | NA |  |
| *Menidia menidia* | Silverside | n=1 |  | NA |  |
| *Coryphaena hippurus* | Dolphinfish | n=1 |  | NA |  |
| *Thunnus thynnus* | Bluefin tuna | n=1 |  | NA |  |
| *Oncorhynchus kisutch* | Coho salmon | n=1 |  | NA |  |
| *Balaenoptera musculus* | Blue whale | n=3 | NA |  |  |
| *Balaenoptera physalus* | Fin whale | n=3 | NA |  |  |
| *Balaenoptera acutorostrata* | Minke whale | n=3 | 25.2, 22.7, 20.4 |  |  |
| *Eschrichtius robustus* | grey whale | n=3 | 24.3, 36.7, NA |  |  |
| *Balaena mysticetus* | Bowhead whale | n=1 | NA |  |  |
| *Delphinus capensis* | Long-beaked dolphin | n=3 | NA |  |  |
| *Delphinus delphis* | Short-beaked dolphin | n=3 | NA |  |  |
| *Phoca vitulina* | Harbor seal | n=3 | NA |  |  |
| *Zalophus californianus* | California sea lion | n=3 | NA |  |  |
| *Carcharodon carcharias* | White shark | n=1 | NA |  |  |

Although *in silico* testing suggested we might see cross-amplification between the humpback whale assay and bowhead whale during *in vitro* testing, the DNA from bowhead whale did not amplify. DNA from blue whale and fin whale also did not amplify. DNA from minke whale amplified (3/3 individuals) and grey whale also amplified (2/3 individuals) (Table 4, Fig 2). Given these results, the specificity of the assay was 81% (Table 3). The C_q_ values for minke and grey whale were, on average, 22.7 and 30.5 (using just two reported C_q_ values). For comparison, the C_q_ for 1 ng/μL of humpback whale DNA is ∼24 (Fig 2). The *in vitro* specificity testing included a standard curve constructed using extracted DNA from one individual (C551) and the efficiency was 92% with a LOQ of 0.1 pg/μL of DNA extract (Fig 2, Panel A, black crosses).

**Fig 2.**
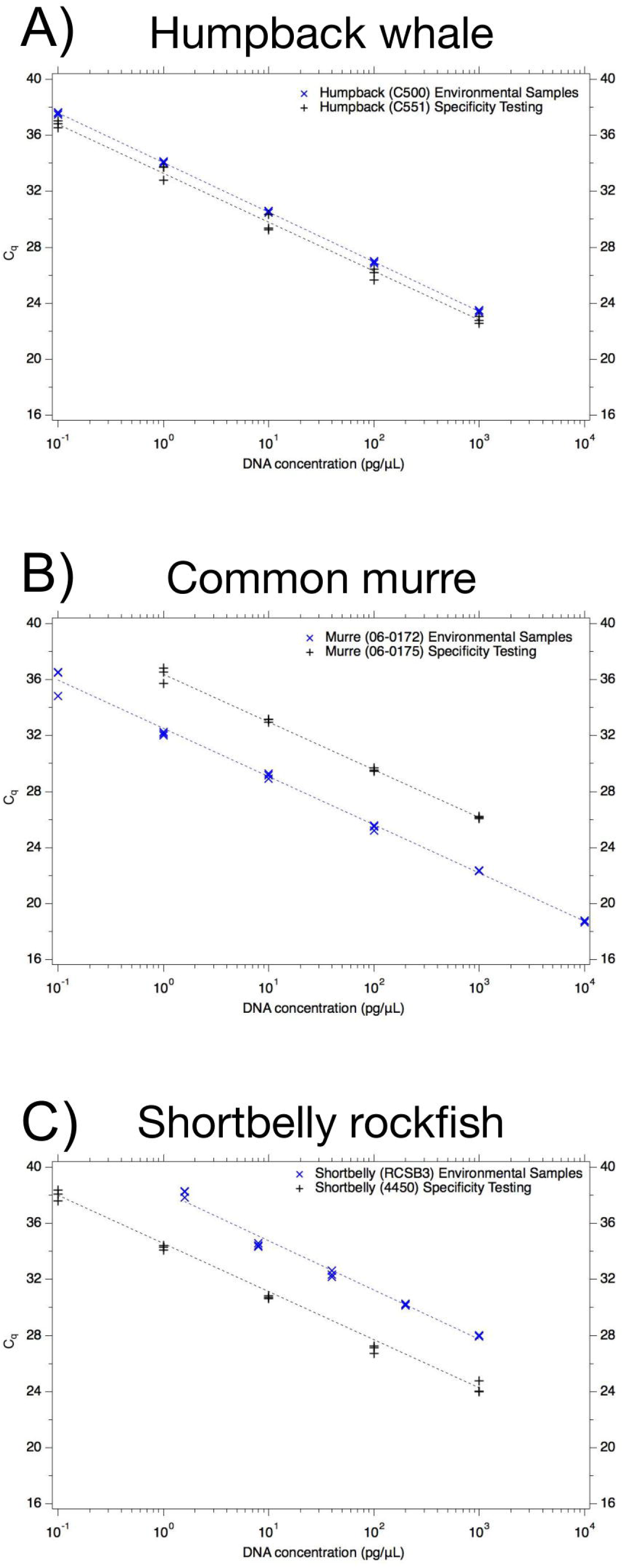
Performance of each assay. Cycle quantification threshold (C_q_) is shown on the y-axis and the x-axis is the concentration of DNA (pg/μL) in the reaction; 2 μL of template was added to each reaction for the dilution series. Panel A shows humpback whale, panel B shows common murre, and panel C shows shortbelly rockfish. For each plot, black crosses are the results of the individuals used for specificity testing (C551 for humpback whale, 06-0175 for common murre, 4450 for shortbelly rockfish). Blue x’s are the results of the individuals used for environmental sample testing (C500 for humpback whale, 06-0172 for common murre, RCSB3 for shortbelly rockfish). In the case of shortbelly rockfish (Panel C), some of the specificity testing was also performed using the standard curve used for environmental sample testing. Each concentration was run in triplicate reactions and in some cases, symbols are overlapping.

For the humpback whale field samples, a standard curve was generated using extracted DNA from a different individual than that which was used for specificity testing (individual C500). The efficiency of the assay was 99% and the LOQ was 0.1 pg/μL of DNA extract (Fig 2, Panel A, blue x’s). All three qPCR triplicates of all three field samples were positive. After performing dimensional analysis from DNA mass per reaction to volume of water filtered, the average concentration of humpback whale DNA in the field samples was 0.03 pg/ml of water (Table 5).

**Table 5.**
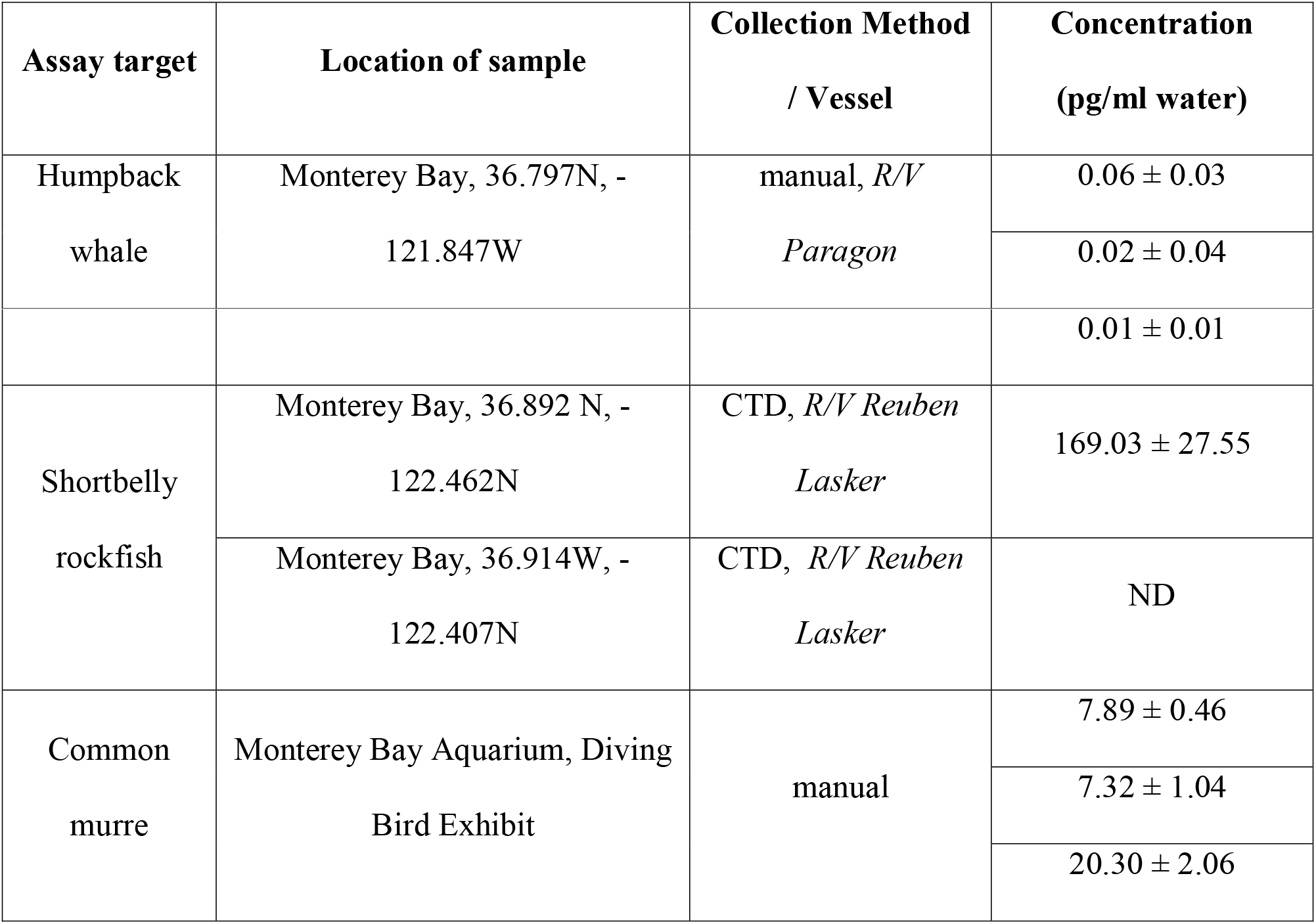
Results of environmental samples tested for each assay. ± is 95% confidence interval. “ND” represents non-detect, meaning sample was not assigned a C_q_ value.

### Shortbelly rockfish assay final design and performance

The shortbelly rockfish assay targets the COI gene. The primers/probe sequences were: F 5’ CAGGAGCATCAGTCGACCTG 3’, R 5’ GAGAAGGAGAAGGACAGCGG 3’, P 5’ FAM - ACACCCTTATTTGTGTGGGCCGTCCT - BHQ 3’ (Table 2). After optimizing primer/probe concentrations and annealing temperature, reactions for amplification were 20 μL reactions with the following: Taqman Universal Mastermix II (1x), forward and reverse primer (0.2 μM), probe (0.1 μM), 2 μL of DNA extract, and molecular-biology-grade water (Sigma-Aldrich, St. Louis, MO). The cycling parameters were as follows: were 95°C for 5 min followed by 40 cycles of 95°C for 15 s, 60°C for 30 s and 72°C for 30 s.

The *in vitro* sensitivity testing demonstrated that all individuals of shortbelly rockfish were positive and therefore assay sensitivity was 100% (Table 3). The logo plots support the sensitivity found *in vitro* as the forward primer, reverse primer, and probe all show unanimous consensus sequences when target sequences were aligned in the regions of the primers/probes (Fig 1, Panel C, top plots).

The assay was specific when tested *in silico* (S2 Table), though the logo plots demonstrate that many of the non-target species within the family have similar sequences to the target sequences (Fig 1, Panel C, bottom plots versus top plots). The plots demonstrate that the probe adds the most specificity with more positions in the probe having different frequencies of base pair occurrence than the target (13/26 base pairs; Fig 1, Panel C, bottom plot). Despite the *in silico* specificity, the specificity testing *in vitro* found that DNA from 2 of 3 individuals of squarespot rockfish (*S. hopkinsi*), and 3 of 3 individuals of both stripetail rockfish (*S. saxicola*) and halfbanded rockfish (*S. semicinctus’)* (Table 4) amplified, all of which also occur in the CCE. Thus, the specificity of the assay was 78% (Table 3). For *in vitro* specificity testing, non-target samples were compared to two standard curves; one was generated using one individual (4450) and had an efficiency of 94% and a LOQ of 0.1 pg/μL (Fig 2, Panel C, black crosses) and the other was generated using DNA extracted from tissue of one target individual of shortbelly rockfish (RCSB3) and had an efficiency of 93% and a LOQ of 1.6 pg/μL (Fig 2, Panel C, blue x’s), All non-target samples that were positive were tested using the RCSB3 standard curve. C_q_ values in the specificity testing of 1-2 ng from these individuals ranged from 23-25 compared to a C_q_ of about 28 for 2 ng (2 μL of 1 ng/μL) of shortbelly rockfish.

Environmental samples were tested to quantify shortbelly rockfish DNA using the standard curve generated using RCSB3 (Fig 2, Panel C, blue x’s). DNA extracted from one of the two environmental water samples did not amplify (ND). The other environmental water sample had an average concentration of 169 pg/mL water (Table 5).

### Common murre assay final design and performance

The common murre assay targets the COI gene. The primers/probe sequences were: F 5’ TGGCGCATGAGCTGGTATAG 3’, R 5’ TATTACAAAGGCGTGGGCGG 3’, P 5’ FAM – ACCGCCCTAAGCCTGCTCATCCGT – BHQ 3’ (Table 2). After optimizing primer/probe concentrations and annealing temperature, reactions for amplification were 20 μL reactions with the following: Taqman Universal Mastermix II (1x), forward and reverse primer (0.2 μM), probe (0.2 μM), 2 μL of DNA extract, and molecular-biology-grade water (Sigma-Aldrich, St. Louis, MO). The cycling parameters were as follows: were 95°C for 5 min followed by 40 cycles of 95°C for 15 s, 64°C for 30 s and 72°C for 30 s.

The *in vitro* sensitivity testing demonstrated that all target individuals were positive, and thus sensitivity was 100% (Table 3, Table 4). The logo plots demonstrate the assay sensitivity by the limited number of base pairs in the primers and probes with multiple frequencies at a specific site (3/20 in forward primer, 0/20 in reverse primer, 1/24 in the probe; Fig 1, Panel B, top plots).

The logo plots for the common murre assay demonstrate the assay specificity by showing the non-target alignment (Fig 1, Panel B, bottom plots) many different base pairs, particularly in the probe (7 of 24, or 30%, of base pairs having different frequencies than the target (Fig 1, Panel B, top plots). Furthermore, the region of the reverse primer was highly conserved and unable to be plotted because no contigs were found when comparing the non-target alignment to the reverse primer sequence (Fig 1, Panel B, bottom plot). The assay was specific both *in silico* (S2 Table, Fig 1) and *in vitro* (i.e., all non-target samples were ND) for the tissue tested (S3 Table, Table 5). Therefore, specificity of the assay is 100% (Table 3). The *in vitro* specificity testing included a standard curve constructed using extracted DNA from one individual (06-0175) and the efficiency was 97% with a LOQ of 1 pg/μL of DNA extract (Fig 2, Panel B, black crosses).

The common murre assay was applied to samples from the Diving Birds Exhibit at the Monterey Bay Aquarium. A standard curve was constructed using a different individual than specificity testing (06-0172). The efficiency was 99% and the LOQ of 0.1 pg/μL of DNA extract (Fig 2, Panel B, blue x’s). The average concentration was 11.8 pg/ml water in the three samples (Table 5).

## Discussion

We developed qPCR assays for three important organisms of the CCE: humpback whale, shortbelly rockfish, and common murre. The assays all had high sensitivity (100% for individuals tested), and the assays yielded positive detection of their targets in environmental samples or samples from an aquarium. However, specificity varied across assays with the common murre assay having high specificity (100%) but the humpback whale and shortbelly rockfish having cross-amplification. The humpback whale assay cross-amplified minke and grey whale both of which also are common in the CCE [18,38]. Thus, positives with the assay might indicate the presence of any one of these whale species. The shortbelly rockfish assay cross-amplified with three species of rockfish also known to be in the CCE. The rockfish genus *Sebastes* includes over 100 species and despite a wide geographical distribution, the phylogeny has been debated amongst researchers for years [23,39]. Therefore, the sub-optimal specificity is not particularly surprising for the rockfish assay. These specificity issues highlight the difficulties in designing qPCR assays, ranging from the availability of sequences in the public databases to biological challenges such as unclear phylogeny and cryptic species.

Design of specific qPCR assays relies upon correct and sufficient genetic data. Public databases such as NCBI’s GenBank contain many entries, however the accuracy and quality of data can be questionable. For example, a recent paper on scyphozoan jellyfishes revealed that the majority of database entries in GenBank for the species *Chyrsaora quinquecirra* are in fact *Chrysaora chesapeakei* [40]. The paper, published in 2017, confirmed the mis-identifications and incorrect entries by both morphological and genetic analyses, yet two years later none of the entries have been corrected in GenBank [40]. Furthermore, processing of Sanger sequencing data that is submitted to public databases requires visual assessment and careful processing and even one base pair difference in a target gene region can impact qPCR assay design and performance. Discrepancies or mistakes in the genetic data used for qPCR assay design can cause poor design and performance and misleading *in silico* specificity testing results. Public databases need better maintenance, a higher standard of metadata required for submission, and accountability for corrections if new evidence suggests that they were incorrect upon submission.

Assuming the genetic data used for assay design are correct, finding regions that are suitable for qPCR chemistry is another challenge for assay design. The targeted region must be relatively short (100-200 bp) in order to capture degraded fragments of eDNA left behind in the water column, but this criterion restricts the possible primer locations on the gene. The chemistry of the qPCR reaction requires certain ranges of melting temperatures, GC content, and other criteria such as the types of base pairs on the 3’ end of the primer in order for the assay to be successful and efficient [29–32]. Due to the narrow range based on the size restriction on the fragment, this can be a major challenge for qPCR assay design. Additionally, the probe, which is often around 25 bp long and thus can be up to 20% of the total target length, must be located in the region between the forward and reverse primers. A probe is included to increase specificity, but it is often difficult to identify a probe that would be successful given the assay’s chemistry requirements. Finally, in assay development there are several trade-offs to be made between assay performance. For example, raising the primer annealing temperature may increase specificity of the assay while simultaneously decreasing sensitivity. Depending on the assay application, these parameters can be optimized ideally with one set of primers and a probe, or many primer/probes can be tested *in vivo*. Furthermore, many studies that publish new qPCR assays provide insufficient detail about methodology, specifically regarding *in silico* and *in vivo* testing. Greater transparency in publications will help guide new efforts for qPCR assay design and testing.

In addition to the technical challenges discussed above, there are several biological considerations when designing an assay. For example, in our study, we designed an assay for shortbelly rockfish. Shortbelly rockfish (*Sebastes jordani)* are one of over 100 species within the genus, which the phylogeny has been and still currently is being debated amongst experts [23,39]. Finding a unique region of a gene for one species in this large genus in the public databases that is of suitable chemistry can be very difficult.

Furthermore, there are differences in individuals from the same species that can produce different qPCR results. We used DNA extracted from tissue samples from different individual target organisms to create standard curves. For humpback whale, the tissue samples yielded similar standard curves when mass of DNA tested is plotted against C_q_ from the qPCR instrument. The common murre and shortbelly rockfish standard curves constructed using DNA from different individuals were different: the same input DNA mass yielded C_q_ values differing by ∼4 C_q_. This suggests differences exist in mtDNA concentrations between the individual DNA extracts. For the two shortbelly rockfish tissue samples, one originated from a fin clip of an individual of unknown age and the other from tissue from the body of a juvenile individual. Both murre tissue samples originated from livers of two individuals of unknown age. The mass of DNA extracted from the various individuals represents contributions from nuclear DNA (nDNA) and mitochondrial DNA (mtDNA). The ratio of nDNA:mtDNA can vary between individuals of a specific species and among tissues in a single individual [41]. A study of their ratio in mice, for example, showed that the ratio varied based on the age (young versus old) of the mouse by a factor of 10 [42]. The ratio seems to be sensitive to DNA extraction method as well [43]. A recent study assessed the differences in eDNA degradation from water samples of a water flea (*Daphnia magna)* when quantified by targeting both a mitochondrial and nuclear gene region [44]. This study found that nuclear DNA was more abundant, however the lengths of the two target regions were also different and the authors acknowledge that other technical biases may be influencing the results such as primer affinity. Different nDNA:mtDNA ratios in the tissue extracts could explain the differences in C_q_ values seen between individuals in our study. Future work should give careful thought to what sort of standard should be used for qPCR assays. Ultimately, this may be informed by what the goal of the measurements are and whether the results need to be externally valid, and relatable to work conducted by others. If so, then synthetic gene fragments may be ideal standards.

The challenge of developing sensitivity and specific qPCR assays is common among a number of fields including that of microbial source tracking, which aims to identify sources of microbial pollutants in the environment using molecular DNA markers, or fragments of DNA unique to a particular organism that indicate its presence. To deal with imperfect qPCR assay sensitivity and specificity, researchers in that field have used Bayesian statistics to interpret qPCR data and provide detection probabilities [45,46]. Bayesian statistics use some elements of prior knowledge to interpret results from molecular assays. For example, in the field of microbial source tracking, authors combine results from multiple qPCR assays, their inferred sensitivity and specificity values, and pre-existing likelihoods of false negatives and positives (Type I and II error rates) from the literature to generate confidence levels to make conclusions using qPCR data [46]. It has been demonstrated that Bayesian statistics can be useful in interpreting eDNA data using model datasets and theoretical frameworks [7,47]. Lahoz-Monfort et al. applied Bayesian techniques to specifically address the issue of false positives errors in eDNA datasets, including using other sampling methods at the same time as eDNA sampling not prone to false positives [47]. Shelton et al. used Bayes’ theorem to develop a framework to estimate species abundance from water samples analyzed for eDNA metabarcoding accounting for all parts of the process from the biomass present at the given sampling location, how the biomass relates to the eDNA concentration, the filtration method, DNA extraction, the stochasticity of PCR amplification, DNA sequencing, and the bioinformatic processes of assigning OTUs to taxa [7]. With the assays presented here, given the less than ideal specificities for the humpback whale and shortbelly rockfish assays, using multiple markers or incorporating other prior knowledge such as visual data (e.g., humpback whale sightings) using a Bayesian approach could improve the interpretation of these assays when applied to field samples. Other eDNA studies have used similar approaches in detection of schistosomiasis [48], carp [49] and in assessing the impact of water filtration and PCR amplification from water samples in an aquarium [50].

Biomonitoring for common murre, rockfish, and whales in the CCE is usually carried out using visual surveys. The assays we developed here, along with assays that have been previously developed for krill, sardines, anchovies, and mackerel [36,51] could be used together to conduct biomonitoring of important vertebrate and invertebrate species from diverse trophic levels in the CCE. The murre assay is sensitive and specific and thus reliable to detect the presence of murre eDNA in a water sample. The shortbelly rockfish assay is genus specific, meaning that positive detections indicate the presence of shortbelly rockfish, but the detection could also be from squarespot rockfish, stripetail rockfish or halfbanded rockfish. Finally, positive detections from water samples using our humpback whale assay could indicate the presence of minke or grey whale. These assays are still useful if any other metadata or identification method is used in conjunction with the qPCR assay or if further testing is performed such as Sanger sequencing the qPCR product to confirm the identification of the target DNA.

eDNA qPCR methods can be implemented on archived, filtered water samples and may allow the development of quantitative temporally and spatially intensive biomonitoring data sets. In addition to increased monitoring efforts on specific species, an important emerging use of such data sets in the CCE is understanding larger scale questions such as how global climate change is affecting food webs [52].

## Acknowledgements

We thank Hilary Starks for preparation of the library samples. We thank Kris Walz for obtaining and performing DNA extractions for tissue used in testing. We thank Carlos Garza and Libby Gilbert and the NOAA Southwest Fisheries Science Center, Barbie Halaska and the Marine Mammal Center, Corinne Gibble and the California Department of Fish and Wildlife, and Nate Rice and the Academy of Natural Sciences of Drexel University for providing samples for sensitivity and specificity testing. We thank Sue Lisin and Aimee Greenebaum for providing access and sampling of the Monterey Bay Aquarium.

## Supporting Information

**S1 Table. Accession numbers used to design primers for each target species.**

**S2 Table. Accession numbers of non-target organism used for specificity testing *in silico*.**

**S3 Table. Metadata of tissue samples tested for sensitivity and specificity.**

